# Subiculum Neurons Map the Current Axis of Travel

**DOI:** 10.1101/050641

**Authors:** Jacob M. Olson, Kanyanat Tongprasearth, Douglas A. Nitz

## Abstract

Travel constrained to paths, a common navigational context, demands knowledge of spatial relationships between routes, their components, and their positioning in the larger environment. During traversal of an environment composed of multiple interconnected paths, a subpopulation of subiculum neurons robustly encoded the animal’s current axis of travel. The firing of these axis-tuned neurons peaked bimodally at head orientations approximately 180 degrees apart. Track rotation experiments revealed that axis encoding carried the spatial reference frame of the larger environment as opposed to the track itself. However, axis-tuned activity of the same subpopulation was largely absent during unconstrained movement about a circular arena. Thus, during navigation in a path-rich environment, subpopulations of subiculum neurons encode the animal’s current axis of travel relative to environmental boundaries - providing a powerful mechanism for mapping of specific relationships between routes, route components, and the larger environment.

---

Hippocampal CA1 neurons are well known for the discrete location specificity of their action potential firing during free-foraging within an arena (O’Keefe and Dostrovsky, 1971; Wilson and McNaughton, 1993). However, such location-specific firing is often modulated when constraints govern the animal’s running behavior and available trajectories (McNaughton et al., 1983; Foster et al., 1989). For example, alternate trajectories taken through the same position frequently yield strong in-field firing differences (Markus et al., 1995; Wood et al., 2000; Frank et al., 2000; Ferbinteanu and Shapiro, 2003), and spatially-specific firing fields can expand and/or repeat when the same shaped path is taken through different regions of an environment (Derdikman et al., 2009; Singer et al., 2010; Nitz, 2011). These results suggest that hippocampal subregion CA1 can and does encode additional navigationally relevant spatial relationships when they are present in the environment.

The dorsal subiculum (SUB) subregion of hippocampus may be primed to encode more complex spatial relationships given its status as a major downstream target of CA1 (Amaral et al., 1991; O’Mara, 2005; Witter, 2006). From the relatively few recording studies available for this major output structure of hippocampus, it is clear that some SUB neurons exhibit place-specific firing akin to that observed in CA1 (Sharp and Green, 1994; Kim et al., 2012). Reported differences relative to CA1 include increased generalization of place fields across similar environments, proportional increases of firing field size to match scaling of arena size, and increased numbers and sizes of firing fields (Barnes et al., 1990; Sharp, 1997; Sharp, 1999; Kim et al., 2012). Other reports have described some SUB neurons as boundary vector cells, with spatial tuning reflecting proximity and orientation to arena borders (Lever et al., 2009; Stewart et al., 2014).

Environments with multiple routes and pathways greatly increase the prevalence of task-related spatial relationships; however, whether SUB captures these more complex spatial features remains to be determined. We therefore obtained multiple single neuron recordings in rats traversing six interconnected routes as part of a simple navigational task (figure 1A, supplemental figures 1-2) and examined firing activity as a function of environmental location and head orientation. Four partially overlapping three-turn routes led to four goal sites where pieces of cereal were given as reward. Two non-overlapping two-turn paths returned the animal to the shared beginning of the four three-turn routes. The arrangement ensured that each track section had a characteristic direction and axis of travel (figure 1B).

**Figure 1.**
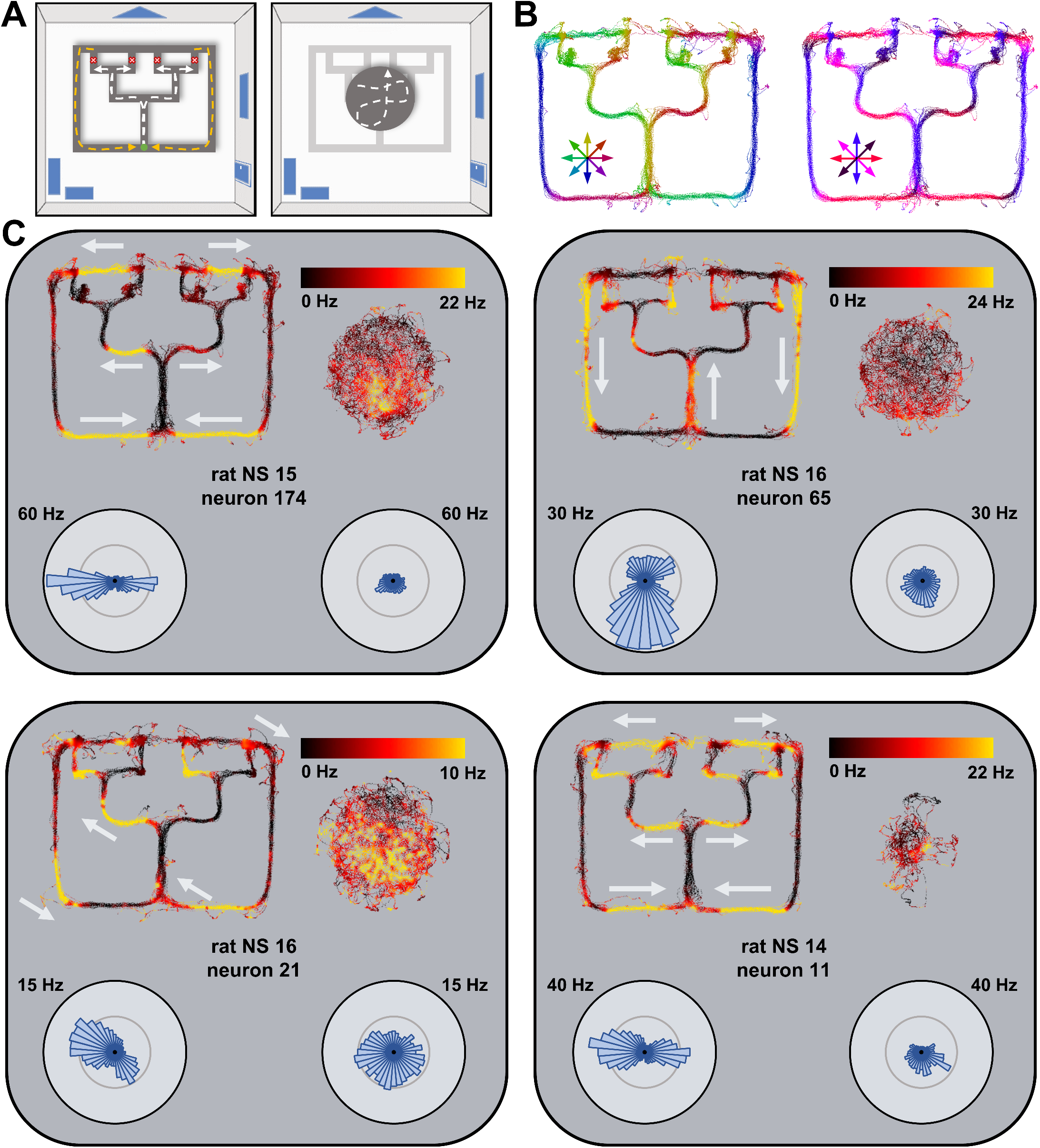
Axis-tuned firing of subicular neurons. **A**. Schematic of route-running and open-field foraging tasks. Left panel depicts the 160 cm X 125 cm track apparatus used in all recordings. *Left panel*: Animals made multiple runs along each of four partially overlapping routes (dashed white lines) leading from a start site (green circle) to any of four goal sites (red circles). From each goal site, the animal returned to the start via either of two return paths (dashed yellow lines). The short (2 cm) edges of the track permitted clear views to the boundaries of the surrounding environment and prominent visual cues (blue shapes). *Right panel*: Recordings were also obtained as animals foraged in a circular, 60 cm-diameter arena also having open view to the surrounding environment. **B**. *Left panel*: Color mapping of the mean directions of travel superimposed on tracking data from a representative recording. *Right panel*: Color mapping of the mean axes of travel for the same animal and recording. This alternate color scheme depicts the axes of travel which are defined by running in either of two opposing directions. **C**. Four examples of axis-tuned SUB neurons from three different animals. Each panel contains an individual neuron’s firing rate color-mapped as a function of track position for all time periods associated with travel >3 cm/second. The directions of travel associated with high firing are indicated by white arrows. For the same neurons, each panel also depicts firing rate as a function of position in the circular arena. Polar plots at the bottom of each panel depict mean firing rate as a function of head orientation during track-running (left) and arena (right) recording epochs. Polar plots are max-normalized to the firing rates of each neuron and labeled according to the firing value of the outside circle’s radius. For each neuron, such directional tuning plots evidence bimodality in tuning peaks with approximate 180-degree separation, consistent with encoding of a single axis of travel. Such tuning was largely absent during the arena foraging task.

Immediately apparent from positional firing rate maps and directional tuning plots was a distinctive subset of neurons that fire strongly whenever the animal ran in either of two opposing directions (figure 1C). Such firing was largely independent of the animal’s position in the room, extending across a wide range of track spaces. Put another way, neurons having this property fire when the animal is travelling in either direction along an axis having a specific orientation.

Because neurons with such axis-specific activity were not reported in prior work utilizing free-foraging in open fields (Sharp and Green, 1994; Sharp, 1997; Sharp, 1999; Brotons-Mas et al., 2010; Stewart et al., 2014), we considered the possibility that axis-specific firing emerges during route-running behavior. We therefore recorded this population of neurons during free-foraging in an open circular arena to determine if axis tuning persists in this well-studied paradigm. The arena was placed just atop and centered upon the track environment with clear view to the same distal landmarks (figure 1A). Consistent with prior work (Sharp and Green, 1994; Sharp, 1997; Sharp, 1999; Brotons-Mas et al., 2010; Kim et al., 2012; Stewart et al., 2014), a subpopulation of SUB cells exhibited spatially-specific firing (supplemental figure 3) akin to that of ‘place cells’ in the hippocampal CA1 subregion. However, neurons with axis tuning on the track exhibited little evidence of axis-tuning firing in the arena (figure 1C). Notably, this result stands in contrast to the behavior of head direction cells in neighboring postsubiculum that exhibit directional tuning in arena environments (Taube et al., 1990). The emergence of axis-specific activity appears to demand the context of route-running.

To assess the prevalence of these axis-tuned neurons in SUB, a quantification of their characteristics is required. Such neurons should exhibit bimodal directional tuning peaks separated by 180 degrees, and firing for each direction should be independent of the animal’s position in the environment. To determine the numbers and orientations of distinct tuning peaks, we fit, for half the data of each neuron, Von Mises mixture models of different orders (supplemental figure 4; supplemental methods). Because each Von Mises distribution in a mixture model can potentially fit an orientation peak, comparison of fits across different model orders can be used to objectively estimate the number of prominent modes in the neural data. Cross-validation of model fitness was carried out on the remaining half of the data. The lowest-order model with a strong fit (>50% decrease in error relative to a 0th-order circular model) and substantial improvement over the preceding model order (a further 20% or more reduction in error) was chosen as the ‘best’ model for each neuron. A higher proportion of SUB neurons were categorized as bimodal (i.e., 2nd-order) than any other model order (single order or 3-8 order; supplemental figure 5) within a wide range of model improvement criteria (10-22.5%). This population bias toward bimodality existed in data collected during track-running but not in arena free-foraging data. Finally, for neurons well fit by a 2nd-order model, applying a threshold for 2X greater firing at model maxima relative to minima removed weakly tuned neurons (figure 2A).

**Figure 2.**
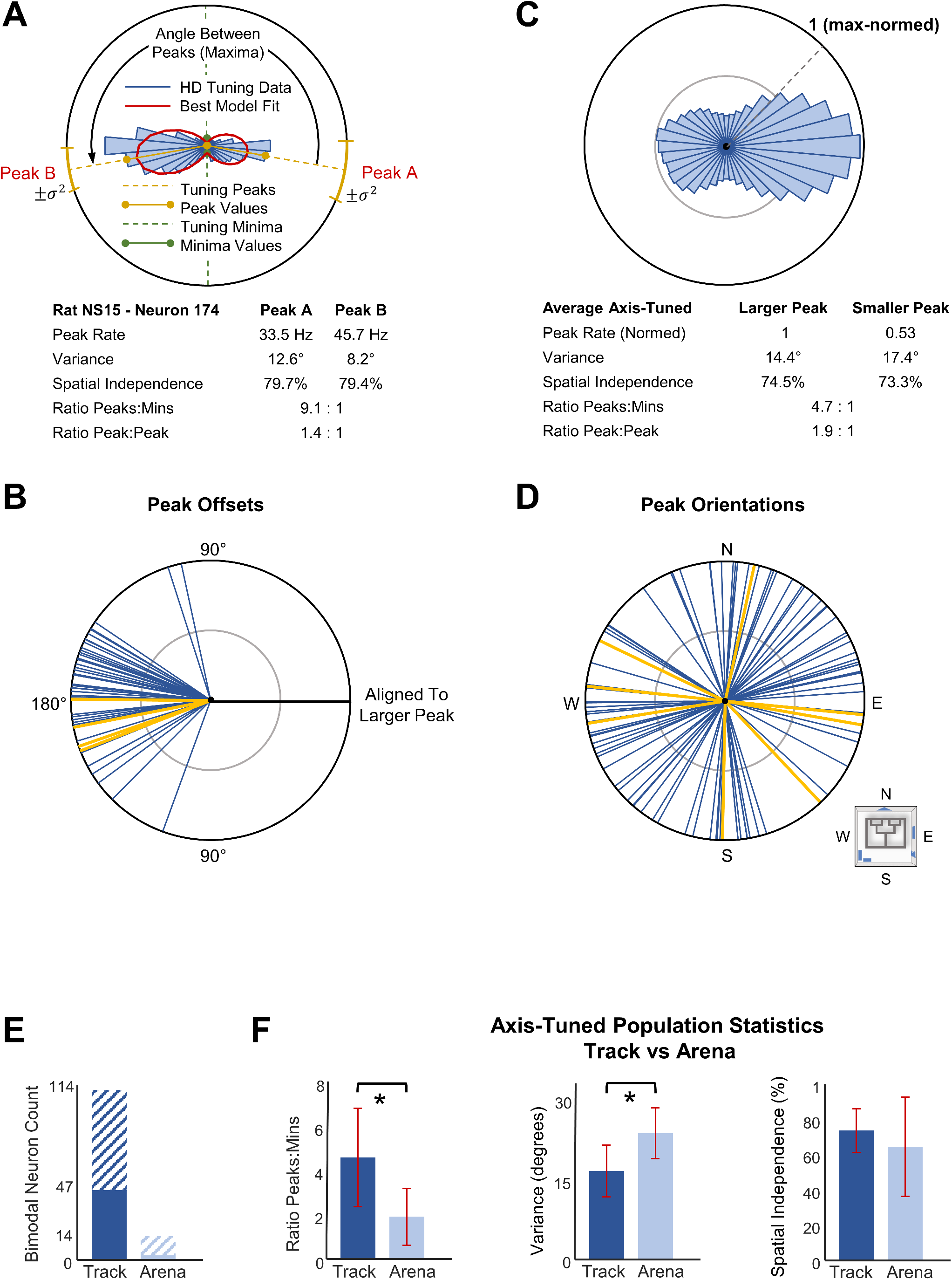
Quantification of axis-tuned firing. **A**. The same directional tuning plot as for the neuron in figure 1C (top left panel) is depicted again to describe measures used to quantify axis-tuned firing. For this neuron, a mixture model using two Von Mises distributions (red ellipses) provided a good fit to the neuron’s directional tuning plot. Models utilizing additional distributions produced marginal increases in fit. The model yields two tuning peaks (maxima) at 100 and 260 degrees from room north (dashed yellow lines) with circular variances shown at the margin (yellow lines). ‘Peak’ firing rates are those of the mean rate for the tuning bins containing the model’s two tuning peaks. Tuning minima are given by the firing rates in tuning bins associated with the model’s tuning minima. These values are used to quantify robustness of tuning (maxima:minima) and to measure bias in tuning for one axis direction versus another (larger peak:smaller peak). Spatial independence scores reflect the percentage of all positions for which directional firing was greater than at least 50% of the mean directional rates for each model-identified peak. **B**. For all 47 SUB neurons meeting criteria for strong axis tuning, the directional tuning plots were aligned to the larger model peak of each neuron (black line). Blue lines depict the relative orientations of these neurons’ opposite, smaller tuning peaks. Nearly all such second peaks fall within 45 degrees of 180-degree separation. Gold lines depict the second-peak orientations for the neurons of figure 1C. **C**. Mean of the max-normalized directional tuning plots for the 47 SUB neurons meeting criteria for strong axis tuning. Tuning plots were first aligned by the bin yielding the highest mean rate. **D**. Orientations of all primary and secondary peaks relative to the space of the surrounding environment. The orientations for the neurons of figure 1C are again shown in gold. **E**. Hashed bars depict the number of SUB neurons (of 542) for which the 2nd-order Von Mises mixture model produced the best fit for track versus arena data. Dark blue and light blue bars depict the number of neurons that also met criteria for ratio of directional tuning maxima versus minima and spatial independence (only 1 of 542 neurons met all criteria for the arena session). **F**. For the 47 neurons meeting criteria for strong axis tuning on the track, peak to minima ratios were significantly higher (left panel; p = 1.79Xe^−11^) for the track recording epoch versus the arena recording epoch, whereas circular variance was reduced (middle panel; p = 2.35Xe^−18^). Spatial independence of directional tuning was statistically similar (p = 0.07; right panel).

We also assessed the positional independence of bidirectional firing to identify those neurons consistently encoding travel along a specific axis. For all track locations associated with either of the neuron’s preferred tuning directions, we determined whether the neuron’s firing rate exceeded at least 50% of the mean rate for those same directions. Neurons were considered spatially independent if the majority of associated locations met this criterion.

The 47 neurons (of 542 tested) meeting these criteria were strongly tuned to a specific axis of travel on the track. Orientation separations between each neuron’s two peaks overwhelmingly clustered near 180 degrees (figure 2B). For each peak, the tuning specificity was high, having a mean circular variance of 0.27 radians (+/- 0.08 STD). Firing rates at the two model-identified tuning peaks (i.e., maxima) were, on average, 4.65X that of firing rates at the minima (+/- 2.23 STD, outlier value of 90.4 removed; figure 2C). Thus, the actual strength in bimodal tuning was well above the applied criterion. Spatial independence of directionally tuned firing was also well above the applied criterion with neurons firing over a mean of 73.9% (+/- 12.5 STD) of all track positions associated with the preferred tuning directions. Across this subpopulation, the preferred tuning directions of firing were evenly distributed (Hodges-Ajne uniformity test, N = 94, p = 0.9182; figure 2D), a result somewhat surprising considering that the track behavior is dominated by travel along the north-south and east-west axes. Taken together, these criteria successfully identify strong examples of axis-specific firing for a significant population of SUB neurons.

By the very same criteria, no neurons with strong axis-tuned firing were found in arena recordings. Furthermore, the 47 neurons identified as having strong tuning during track-running exhibited limited evidence for axis tuning in the arena but instead a range of spatial tuning consistent with that previously reported (Sharp and Green, 1994; Sharp, 1997; Sharp, 1999; figures 2E, 2F, supplemental figure 6). The 2nd-order model proved best for only 1 of the 47. In fact, the 2nd-order model proved best for only 14 of the entire sample of 542 neurons. Relative to measures obtained for track running epochs, model maxima to minima ratios of track axis-tuned neurons were significantly reduced (N = 47, p = 1.79Xe^−11^, Wilcoxon rank sum test). Circular variance in peak tuning was increased for these same neurons in arena recordings (N = 94, p = 2.35Xe^−18^, Wilcoxon rank sum test). Spatial independence was unchanged across conditions (N = 94, p = 0.07, Wilcoxon rank sum test), a result simultaneously consistent with axis-tuned firing on the track and the broad spatial tuning typical of SUB arena data (Barnes et al., 1990; Sharp and Green, 1994; Kim et al., 2012; supplemental figure 6). Thus, axis-tuned firing during movement in the open arena was greatly impoverished relative to track running epochs.

The preceding results strongly indicate that the context of track-running and/or the constraints on available trajectories engendered by the track apparatus itself are critical in generating axis-tuned firing of SUB neurons. As the structure of the track allowed clear view to distal visual cues along the walls of the recording room, axis-tuned neurons could, in principle, take either the recording room or the track structure itself as the spatial frame of reference. To determine which frame was responsible for organizing such activity, a partially overlapping subset of neurons (N = 170) were also recorded following 90-degree rotations of the track relative to the room (figure 3A). Under such conditions, neurons meeting the aforementioned criteria for strong and spatially reliable axis-tuned firing in either of the two track orientations (N = 26) remained anchored to the spatial frame of reference given by the recording room. Highly similar directional tuning plots were observed for the standard and rotated conditions, consistent with the fixed position of the tracking camera relative to the room (figure 3B). Correlations between directional tuning curves for the standard and rotated track configurations had a mean of 0.76 (+/- 0.16 STD). When the directional tuning curve for the rotated track session was rotated to maintain the relationship to the track, mean correlations between the two sessions were significantly weaker (mean r = −0.41 +/- 0.22 STD; N = 26, p = 6.55Xe^−10^, Wilcoxon rank sum test; figure 3C). This result confirms statistically that axis-tuned firing was anchored to the room as opposed to the track.

**Figure 3.**
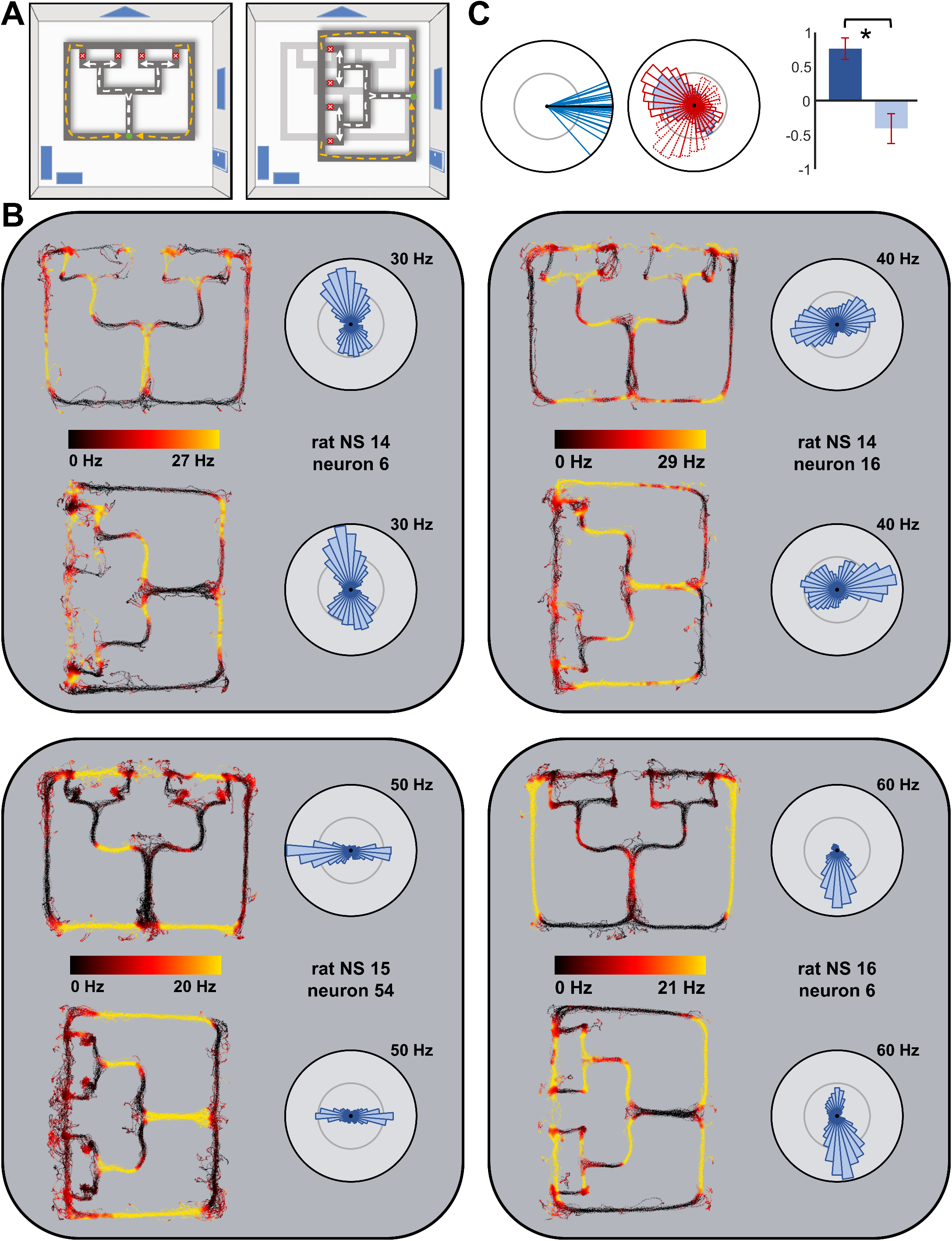
Determination of spatial frame of reference for axis tuning. **A**. Schematic depicting the normal (left panel) and rotated (right panel) track placements. **B**. Color-mapped firing rates for axis-tuned neurons in the normal and rotated track configurations alongside the associated directional tuning plots. The neuron in the lower left panel is repeated from figure 1C (upper left panel). In all four cases, the tuning peak orientations are largely unchanged across conditions indicating that the surrounding environment, as opposed to the track itself, serves as the spatial frame of reference. Because of this, the track positions yielding maximal firing in the rotated condition differ substantially from those for the normal track orientation. **C**. Comparison of orientation tuning across track orientations. *Left panel*: Black line indicates the alignment of the largest tuning peak on the normal track alignment for all 26 neurons meeting axis-tuning criteria in the track/rotation experiment subpopulation. Blue lines show orientations of the nearest peak from the 90-degree track rotation data. *Center panel:* To determine spatial frame of reference, correlations of the directional firing rate values for the normal track condition (blue) and rotated track condition (red) were compared to correlations between track condition (blue) and 90-degree rotation of the rotated track data (dotted red). *Right panel:* Mean (+/- STD; N = 26) of the correlations between directional firing rate values for the normal and rotated track conditions is shown in dark blue. Light blue bar depicts the mean of correlations for the same data following 90-degree rotation of the data from the rotated track session (matching track rotation; p = 6.55Xe^−10^).

Taken together, the present analyses of directional and spatial tuning properties of SUB neurons reveal a previously unknown form of orientation encoding, the animal’s current axis of travel. Axis-tuned firing in a subpopulation (approximately 9%) of SUB neurons is recognized and quantified from directional tuning plots characterized by two distinct peaks in firing rate, separated by approximately 180 degrees. Such axis-tuned firing: 1) can exhibit near complete independence from environmental location; 2) is expressed primarily in the context of route-running along tracks as opposed to unconstrained movement in an open arena; and 3) carries the boundaries of the larger environment (a.k.a., allocentric space) as its spatial frame of reference. Notably, the firing patterns of SUB axis-tuned neurons are clearly distinguished from those of other known forms of spatial information encoding, including hippocampal and SUB place cells (O’Keefe and Dostrovsky, 1971; Wilson and McNaughton, 1993; Sharp and Green, 1994), medial entorhinal cortex (MEC) grid cells (Hafting et al., 2005), MEC border cells (Savelli et al., 2008; Solstad et al., 2008), and even head direction cells (Taube et al., 1990), whose activity maps only a single orientation and whose tuning is strongly present during free-foraging in open arenas. Previous work has described the firing of some SUB neurons according to a boundary vector model (BVM). In this model, spatial tuning is determined by the proximity and orientation of a border to the animal (Barry et al., 2006; Lever et al., 2009; Stewart et al., 2014). Considering all track edges as the relevant boundaries, the axis tuning of SUB neurons seen in the present work is, on the one hand, largely consistent with such properties and may reflect unique aspects of their expression in track-based environments. On the other hand, axis-tuned neurons on the track robustly maintained their preferred orientations following track rotations but did not exhibit the BVM-predicted annular shaped firing fields in the arena (supplemental figure 6).

The context dependency of axis-tuned SUB activity parallels that of several other forms of spatial representation. For hippocampal neurons, route-running is associated with directional dependence in their place-specific firing (McNaughton et al., 1983; Markus et al., 1995), the emergence of trajectory-specific place fields (Wood et al., 2000; Frank et al., 2000; Ferbinteanu and Shapiro, 2003), and the emergence of repeating spatially-specific firing patterns for paths having equivalent shapes (Singer et al., 2010; Nitz, 2011). The presence of high walls defining a ‘switchback’ path can result in environmental fragmentation and associated resetting of MEC grid cell alignment across repeating route subspaces (Derdikman et al., 2009). Finally, the action correlates of most posterior parietal cortex neurons in free-foraging are replaced by route-position-dependent firing on tracks (Nitz, 2006; Nitz, 2012; Whitlock et al., 2012). Thus, route-running induces qualitative changes in spatial mapping in several brain structures and such changes yield conjunctive information concerning relationships between paths and the space of the larger environment (see also, Alexander and Nitz, 2015). The near absence of a route-running context in prior SUB recording experiments may well explain why axis-tuned firing has not previously been detected.

The expression of axis-tuned SUB activity within an environment composed of multiple interconnected paths is perhaps an initial clue to its functional significance. Axis-tuned firing essentially represents travel along a line having a particular orientation relative to the environment. As such, a potential function of axis-tuned activity could be to encode the orientation of track segments relative to the larger environment. Such a function could be particularly relevant in real-world urban or rural environments where, in contrast to laboratory-based high-walled arenas, environmental boundaries can be difficult to precisely define and visual access to boundary landmarks may vary considerably. For example, a commonly used orientation tool in human navigation is to align oneself to a city’s street grid based on a prominent, well-known, linear landmark such as a coastline, river, or well-recognized street. At the level of systems neurophysiology, this could be accomplished by pairing of axis-tuned activity with the place specific firing of hippocampal and SUB neurons. Together, these forms of spatial information could generate a robust mapping of path position and orientation relative to environmental position. Such a process may well explain the presence of conjunctive representations of route and environmental position in the firing of neurons in retrosplenial cortex (Alexander and Nitz, 2015).

Finally, layering representation of movement axes on top of the cognitive map provided by hippocampal place cells and MEC grid cells yields a more functionally complete mapping of space. During travel along routes, it is likely relevant that any single axis-tuned neuron will be iteratively co-active with a specific subset of all hippocampal neurons exhibiting space and direction-specific firing fields (i.e., place fields). Assuming connectivity with synaptic plasticity between axis-tuned neurons and hippocampal place cells, it follows that activation of similarly axis-tuned SUB neurons could specify the subset of place cells whose firing fields are consistent with plausible trajectories from the current position as well as the positioning of alternate pathways that allow movement along the same orientation. In this way, axis-tuned neurons could positively bias the cognitive map toward the generation of the planned trajectories known to occur during hippocampal sharp-wave/ripple events (Foster and Wilson, 2006; Diba and Buzsaki, 2007; Gupta et al., 2010; Jadhav et al., 2012; Pfeiffer and Foster, 2013).

---

**Supplemental Figure 1**. Summary of recording site histological data. Recordings of subiculum neurons (N = 542) were obtained from a total of five four-tetrode bundles in three animals. Numbers of total recorded neurons and numbers of neurons with axis-tuned firing are included above each figure. Red arrows depict tracks left by the bundles and their approximate endpoints. Three of the recording sites were restricted to the subiculum, and two (NS15-left and the lateral bundle in NS16-right) were in a transition zone bordering the CA1 subregion. Abbreviations: RCTX (retrosplenial cortex), DG (dentate gyrus), SUB (subiculum).

**Supplemental Figure 2**. Waveform discrimination. **A**. Waveform discrimination for five axis-tuned neurons recorded on the same tetrode from one recording of rat NS14. *Center:* Color-coded clusters of individual waveforms in a 3D plot of peak-valley voltage on wires 2, 3, and 4 (wire 1 not recorded). For each cluster, the color-coded waveform plots of all waveforms were included. *Surrounding polar plots:* Adjacent polar plots to each waveform plot show the directional tuning of that neuron on the track. **B**. Waveform discrimination of one axis-tuned neuron (red) with all other waveforms recorded (black points) shown from rat NS15. The same three plots are included here as in panel A, but the peak-valley voltage plot is only of wires 1 and 2. Wires 3 and 4 were not recorded on this tetrode. **C**. Waveform discrimination for three neurons recorded on a single tetrode from rat NS16. Here, two are axis-tuned and one is not.

**Supplemental Figure 3. A**. Neurons (N = 542) plotted based on spike width versus burst index. The distribution is consistent with prior work (Kim et al., 2012). **B**. Similar to previous studies, the degree of spatially-specific activity among the population varied greatly. Shown are example neurons whose positional firing rate maps reflect the observed range of low (top panels) to high specificity as measured by information-per-spike (Skaggs et al., 1993). Note that the top right and bottom right panels correspond to neurons found to have robust axis-tuned firing during track-running.

**Supplemental Figure 4**. Model Schematic. **A**. Example neuron training data and model fits. For all plots, directional tuning data is shown from a randomly-selected half of the track data used for training (blue rose plots). Von Mises mixture model fits of each order used (0-8) are overlaid (red ellispses). **B**. Directional tuning plots from the remaining half of data were used for cross-validation. Sum-squared-error (SSE) between the cross-validation data and each order model (red, same models as in A) is printed above each figure along with its SSE Ratio, the remaining error normalized by the amount of SSE of the naïve circular model (order 0). For a model to be considered a ‘best fit’, the SSE Ratio must be below 0.5 (<50% error of naïve circular model). Among model orders with SSE Ratios below 0.5, the difference between the model and preceding order model must be greater than 0.2 (20% improvement; difference value printed above plots, see also supplemental figure 5). For this neuron, the criteria lead to selection of the 2nd-order model (red box). C. To be considered as strongly axis-tuned, two more criteria must be met. First, the ratio of mean actual data firing rate at model maxima to mean actual data firing rate at model minima must exceed 2. Mean peak (maxima) values for this neuron (green lines) were 8.2X mean minima values (black lines). Finally, the spatial independence of each model’s maxima must both exceed 50%. For all track locations associated with movement in either of the two preferred directions, the neuron was considered ‘active’ if its mean rate at that position and orientation was at least 50% of the overall mean rate for that direction. In the case of this neuron, most of the points in the light green orientation lie along the light green arrows on the right panel (see also figure 1B – right panel) and the points in the dark green orientation lie along the dark green arrows. Neurons were considered spatially independent if the majority of associated locations met this criterion. For this sample neuron, the larger, light green peak had 88% spatial independence and the smaller, dark green peak had 58% spatial independence. Because it met all three criteria, this neuron is in the axis-tuned subpopulation.

**Supplemental Figure 5**. Model Parameter Flexibility. To select the best order for the Von Mises mixture model for each neuron, we trade off model complexity with fit improvement by selecting the most complicated model that yields 20% improvement in sum squared error over the preceding order model. The criterion is arbitrary but qualitatively consistent across a wide range of values. Here we plot the number of neurons in the population categorized into each order type based on a range of criterion values. Criterion values ranged from 2.5-40% by increments of 2.5% (blue lines, dark colors – light colors = low to high criterion values). We selected the 20% value for our criteria (red lines). Track model fitting data is given in the left panel. From 10-22.5%, more neurons are classified as 2nd-order than any other order. This demonstrates that this is a property of the population and not the model parameters, especially when compared to platform data (right panel), where no model parameter results in a population bias to 2nd-order (bimodal) mixture models.

**Supplemental Figure 6**. For each neuron meeting criteria for strong axis-tuned firing during the track running session, the positional firing map for the same neuron is shown for the arena foraging session. Peak (arena max) firing rates are given above each.

## Supplemental Methods

### Subjects

All subjects were male Sprague-Dawley rats (N = 3). Rats were housed individually and kept on a 12-hour light/dark cycle. Prior to experimentation, animals were habituated to the colony room and handled for 1-2 weeks. During training and experimentation, rats were food restricted and weights were maintained at 85%-95% of free-fed weight with water available continuously. Rats were required to reach a minimum weight of 350 g (5-10 months of age) prior to surgery and subsequent experimentation. All experimental protocols adhered to AALAC guidelines and were approved by IACUC and the UCSD Animal Care Program.

### Apparatus

Behavioral tasks were conducted using both a circular wall-less arena and a triple ‘T’ track maze. The track (figure 1A, left panel, 8cm-wide pathways, overall 1.6m × 1.25m in length and width, painted black) stood 20 cm high in the middle of a large recording room and was visually open to prominent distal cues. The track edges were only 2cm in height, allowing for an unobstructed view of the environment. The arena (figure 1A, right panel, 60 cm in diameter) was placed 20 cm above the center of the track. The arena was also visually open to the same prominent distal cues as well as the track below. For the first recording of rat NS14 (N = 17 neurons), a high-walled pot (30 cm in diameter, 22 cm walls) was used in place of the arena.

### Behavior

Rats were habituated to the maze during two 30-minute periods of free exploration. Animals were then trained to run ballistically from the midpoint of one of the long edges of the maze into the center of the apparatus and continue until reaching the long edge opposite the start point (figure 1A, white dashed lines). This consisted of straight sections interleaved with three left or right turns for a complete path run. The total path lengths were 140cm, with turns at 51 cm, 87 cm, and 118 cm. Reward (1/2 piece of Cheerio’s cereal) was made available at the four reward sites. Over 1-2 weeks, animals were trained by approximation to make route traversals between food reward sites. Over another 1-2 weeks, animals were trained by simple trial and error to a criterion of 80% for ballistic (uninterrupted) path traversal. Once animals met criterion, they were trained 2-3 times on the track in the normal orientation, immediately followed by training on the track in the 90-degree rotated orientation. This established familiarity with the rotated track, but the rats were not extensively trained in this orientation.

Multiple reward paradigms were used across the set of animals. In an all-but-repeats paradigm, used for animal NS14, the animal was rewarded at any of the four locations except when the animal repeated the same location as the previous run. In a visit-all paradigm, the animal was rewarded at all locations, but needed to visit all locations before rewards were reset at all paths.

In the arena, two different behavioral epochs were used, each for approximately half of the time in the arena. For the first half of the time, the animal was cued to make trajectories across the full platform for a 1/4 Cheerio’s cereal reward at the track edge. The trajectory orientation was varied in order to obtain adequate sampling of the full platform surface. This pattern produced running activity similar to the track apparatus. The second half of each arena session was free-foraging for small pieces of Cheerio’s reward dispersed randomly in the arena. Data was analyzed together in all portions of the paper to maximize sampling.

### Surgery

Rats were surgically implanted with tetrode arrays (twisted sets of four 12.5-µm nichrome wires) inserted into custom-built microdrives (4-8 tetrodes per microdrive). Rats were implanted bilaterally with 2-3 microdrives into dorsal subiculum. Rats were anesthetized with isoflurane and positioned in a stereotaxic device (Kopf Instruments). Following craniotomy and resection of dura mater, microdrives were implanted relative to bregma (A/P −5.6 – −6.6 mm, M/L ±1.6 – ±2.7 mm, D/V −1.5 – −2.2 mm).

### Recordings

After recovery from surgery, animals were trained for at least one week to ensure adequate behavior and running ability with the new weight of the implant. Electrodes were moved ventrally in 40 µm increments between recordings to maximize the amount of distinct units collected. Each microdrive had 1-2 electrical interface boards (EIB-16, Neuralynx) connected to a single, amplifying headstage (20X, Triangle Biosystems). A tether led to a set of preamplifiers (50X) and a high pass filter (>150Hz). Signals then fed into the acquisition computer running Plexon SortClient software, filtered at 0.45 – 9 kHz, further amplified 1 – 15X (to reach a total of 1,000 – 15,000X), and digitized at 40 kHz. Single units were isolated in Plexon OfflineSorter software. Waveform parameters used were peak height, peak-valley, energy, full-width half-maximum, and principal components. Waveform clusters appearing to overlap with the amplitude threshold set for collection were discarded to avoid collection of neurons with partial spiking data. Waveform amplitudes were monitored to ensure systematic fluctuation did not produce confounds in isolating single units.

Animals’ position was tracked using a camera set 8.5 ft above the recording room floor. Plexon CinePlex Studio software was used to detect red and blue LED lights placed on the animal’s surgical implant, centered on the animal’s head and separated by approximately 5 cm. Position location of the lights was captured at 60 Hz. The animal’s position and orientation was determined by averaging the location of the two lights and calculating the orientation of the vector between the lights. Using the fact that the track apparatus was squared to the room, we averaged the orientation of all time periods with >3 cm/second running and positions on the middle half of the return arms of the track. This angle was defined as 0 degrees, or “room north,” for the recording and was used to align the animal’s heading to the room. Recordings lasted approximately 45 minutes for arena and track recordings and 1 hour 15 minutes for recording sessions with track rotation data. The animal would run in the arena for 3-10 minutes and then on the track for approximately 80 rewarded runs (figure 1A). For track rotation recordings, the animal had access to water for 5 minutes after completing the first session while the track was wiped down and rotated and then ran for another 80 rewarded runs (figure 3A).

We recorded a total of 542 SUB neurons across three rats (N = 81/321/140). For the first animal, all 81 neurons were recorded from the right hemisphere. For the second animal, NS15, 127 neurons were recorded from left SUB and 194 from right SUB. For the third animal, 42 and 98 neurons, respectively, were obtained from medial and lateral tetrode bundles in the right hemisphere. No neurons were excluded from analysis, even if activity was minimal.

### Histology

Animals were perfused with 4% paraformaldehyde (vol/vol) under deep anesthesia. Brains were removed and sliced into 50-µm sections and Nissl-stained to reveal the final depth of electrode wires in SUB. Microdrive depth monitored across recordings and final electrode depth as observed in histology were compatible in all cases.

### Positional Firing Rate Maps

To characterize the firing activity of the SUB neurons, we calculated their positional firing rates by dividing the total number of spikes at each location by the total occupancy time at each location. To include only data where the animal is running, we excluded all samples with less than 3 cm/second velocity or greater than 20 radians/second angular velocity. The latter threshold was used to exclude cases of rapid head turning in the absence of locomotion. Positional firing maps were smoothed using a 2D convolution with a Gaussian filter with s.d. of 2 cm that also accounts for bins with no occupancy. Data were downsampled to 2 cm × 2 cm bins (not smoothed) for analysis of spatial independence of directional firing. For supplemental figure 6, data is downsampled to 2 cm × 2 cm bins and smoothed using a 2D convolution with a Gaussian filter with s.d. of 4 cm.

### Directional Tuning Vectors

Head direction tuning vectors were calculated using the same sample of running data as the positional firing rate maps (i.e., using the same velocity thresholds). Head orientations were binned into 36 10-degree bins. The total number of spikes per bin was divided by the total time in each bin to calculate the mean directional firing rate.

### Orientation Maps

Orientation maps (figure 1B) were created by calculating the mean circular direction (Berens Circular Statistics Toolbox in Matlab) of all samples for a given 2 cm by 2 cm spatial bin.

### Burst Index for Spiking Activity

To assess burstiness in SUB spiking activity, we applied the method outlined and utilized by Kim et al. (2012). We did this for two reasons. First, the method developed by Kim et al. is not confounded by firing rate differences, a key factor considering the diverse firing rates of SUB neurons. Second, using the same method allowed direct comparisons of our results to previous findings. The burst index is computed by integrating the spike autocorrelogram from 1-6 ms and dividing the result by the integrated power from 1-20 ms. The measure allowed us to demonstrate that the basic firing properties of SUB neurons recorded in our work are in line with those observed previously (Kim et al., 2012).

### Fitting of Von Mises Mixture Models

To fit direction tuning data, we utilized Von Mises distributions, a periodic generalization of the Gaussian distribution. Because our data potentially displayed multiple peaks, different orders of von Mises mixture models were fit and compared. The 0th-order model is equivalent to a uniform circular distribution whose values, like those of the normalized rate data, sum to 1. Order 1 to order 8 models employ 1-8 von Mises distributions. For each condition and neuron, a directional tuning vector was calculated from a randomly-selected half of the data. Mean firing rates were converted to proportional data samples in the mean direction for each bin. Then, for each order model from 1-8, we implemented the estimation-maximization (EM) algorithm (Dempster et al., 1977) to find a model fit to the data. Initialization used Von Mises distributions equally spaced around the circle with ten different mean starting points that uniformly sampled the possible angle space. The stopping criterion for the algorithm was an improvement of less than 0.01 percent in the log likelihood value, and the best fitting model of each order as determined by log likelihood was the model used for that order.

### Evaluation of Von Mises Mixture Model Fit

For each neuron, each model was cross-validated by calculating the sum-squared-error (SSE) between model values and actual direction tuning values for the remaining half of the data. The ratio of SSE of each order mixture model to SSE of the circular (0th-order) model for that same neuron was used to determine overall goodness of fit. Taking into account the trade-off between model fit and model complexity, we defined the ‘best’ model as the model yielding a 50% improvement in fit over the circular model and a 20% improvement over the next simplest model. Similar results were obtained using thresholds of 40-60% improvement over the circular model and 10-22.5% improvement over the next simplest model (supplemental figures 4, 5)

### Spatial Independence

The spatial independence criterion was used to identify neurons that were consistently active when the animal was facing a preferred orientation regardless of spatial location on the track. For all 2 cm by 2 cm spatial bin locations with at least 10 samples in a peak 10-degree bin orientation, the neuron was considered active if its mean rate at that position and orientation was at least 50% of its overall mean rate for that orientation. If more than 50% of the viable locations for both peaks of a neuron were active, the neuron was considered to meet this criterion.

### Maxima and Minima Orientations and Ratios

Peak orientations are the Von Mises mixture model orientation parameters. Large and small peaks are determined by the mixture parameters of the model. Maxima and minima values for ratio calculations are determined by actual data values from the orientation bins containing the model maxima and minima. Peak to minimum ratios are the mean of the peak values divided by the mean of the minima values. Peak to peak ratios are simply the larger peak value divided by the smaller peak value.

### Correlation Across Track Positions

Pearson correlations for the track rotation experiment were calculated between the directional tuning vectors of the track data in the normal orientation and directional tuning data from the 90-degree rotated track session for each individual neuron. Pearson correlations were also calculated between the normal track orientation data and the 90-degree rotated track data shifted 90 degrees. A Wilcoxon ranked sum test was run between the two populations for the axis-tuned subpopulation.

### Alignment of Peaks

Peak alignments for the track rotation experiment are calculated using the angle difference between the larger peak on the non-rotated track and the closest peak on the rotated track.

## References

Alexander AS, Nitz DA. (2015) Retrosplenial cortex maps the conjunction of internal and external spaces. Nat Neurosci. 18(8):1143–51.

Amaral DG, Dolorfo C, Alvarez-Royo P (1991) Organization of CA1 projections to the subiculum: a PHA-L analysis in the rat. Hippocampus 1:415–436.

Barnes CA, McNaughton BL, Mizumori SJ, Leonard BW, Lin LH.(1990) Comparison of spatial and temporal characteristics of neuronal activity in sequential stages of hippocampal processing. Prog. Brain Res. 83:287–300.

Barry C, Lever C, Hayman R, Hartley T, Burton S, O’Keefe J, Jeffery K, Burgess, N (2006) The boundary vector model of place cell firing and spatial memory.

Brotons-Mas JR, Montejo N, O’Mara SM, Sanchez-Vives MV. (2010) Stability of subicular place fields across multiple light and dark transitions. Eur J Neurosci. 32(4):648–58.

Derdikmanet al. (2009). Fragmentation of grid cell maps in a multicompartment environment. Nature Neuroscience, 12(10), 1325–32.

Dempster, A. P., Laird, N. M., & Rubin, D. B. (1977). Maximum likelihood from incomplete data via the EM algorithm. J R Stat Soc Series B (Methodological). 39(1):1–38.

Diba K, Buzsáki G. (2007) Forward and reverse hippocampal place-cell sequences during ripples. Nat Neurosci. 10(10):1241–2.

Ferbinteanu J, Shapiro ML. (2003) Prospective and retrospective memory coding in the hippocampus. Neuron 40(6):1227–39.

Foster TC, Castro CA, McNaughton BL. (1989) Spatial selectivity of rat hippocampal neurons: dependence on preparedness for movement. Science 244(4912):1580–2.

Foster DJ, Wilson MA. (2006) Reverse replay of behavioural sequences in hippocampal place cells during the awake state. Nature 440(7084):680–3.

Frank LM, Brown EN, Wilson M. (2000) Trajectory encoding in the hippocampus and entorhinal cortex. Neuron 27(1):169–78.

Gupta AS, van der Meer MA, Touretzky DS, Redish AD. (2010) Hippocampal replay is not a simple function of experience. Neuron 65(5):695–705.

Hafting, T., Fyhn, M., Molden, S., Moser, M.-B., & Moser, E. I. (2005). Microstructure of a spatial map in the entorhinal cortex. Nature, 436(7052), 801–6.

Jadhav SP, Kemere C, German PW, Frank LM. (2012) Awake hippocampal sharp-wave ripples support spatial memory. Science 336(6087):1454–8.

Kim SM, Ganguli S, Frank LM. (2012) Spatial information outflow from the hippocampal circuit: distributed spatial coding and phase precession in the subiculum. J Neurosci. 32(34):11539–58.

Lever C, Burton S, Jeewajee A, O’Keefe J, Burgess N (2009) Boundary vector cells in the subiculum of the hippocampal formation. J. Neurosci. 29(31):9771–9.

Markus EJ, Qin YL, Leonard B, Skaggs WE, McNaughton BL, Barnes CA. (1995) Interactions between location and task affect the spatial and directional firing of hippocampal neurons. J Neurosci. 15(11):7079–94.

McNaughton BL, Barnes CA, O’Keefe J. (1983) The contributions of position, direction, and velocity to single unit activity in the hippocampus of freely-moving rats. Exp Brain Res. 1983;52(1):41–9.

Nitz, D. A. (2006) Tracking route progression in the posterior parietal cortex. Neuron, 49(5), 747–56.

Nitz, D. A. (2011) Path shape impacts the extent of CA1 pattern recurrence both within and across environments. Journal of Neurophysiology, 105(4), 1815–24.

Nitz, D. A. (2012) Spaces within spaces: rat parietal cortex neurons register position across three reference frames. Nature Neuroscience, 15(10), 1365–7.

O’Keefe, J., & Dostrovsky, J. (1971) The hippocampus as a spatial map. Preliminary evidence from unit activity in the freely-moving rat. Brain research,34(1), 171–175.

O’Mara S (2005) The subiculum: what it does, what it might do, and what neuroanatomy has yet to tell us. Journal of Anatomy 3:271–82.

Pfeiffer BE, Foster DJ. (2013) Hippocampal place-cell sequences depict future paths to remembered goals. Nature 497(7447):74–9.

Savelli F, Yoganarasimha D, Knierim JJ. (2008) Influence of boundary removal on the spatial representations of the medial entorhinal cortex. Hippocampus 18(12):1270–82.

Sharp PE, Green C (1994) Spatial correlates of firing patterns of single cells in the subiculum of the freely moving rat. J. Neurosci 14(4):2339–2356.

Sharp PE. (1997) Subicular cells generate similar spatial firing patterns in two geometrically and visually distinctive environments: comparison with hippocampal place cells. Behav Brain Res. 85(1):71–92.

Sharp PE. (1999) Subicular place cells expand or contract their spatial firing pattern to fit the size of the environment in an open field but not in the presence of barriers: comparison with hippocampal place cells. Behav Neurosci. 113(4):643–62.

Singer AC, Karlsson MP, Nathe AR, Carr MF, Frank LM. (2010) Experience-dependent development of coordinated hippocampal spatial activity representing the similarity of related locations. J Neurosci. 30(35):11586–604.

Skaggs, W. E., McNaughton, B. L., Gothard, K. M., & Markus, E. J. (1993). An information-theoretic approach to deciphering the hippocampal code. In Hanson SJ, Cowan JD, Giles CL. Adv. Neur. In. 5:1030–37.

Solstad T, Boccara CN, Kropff E, Moser MB, Moser EI. (2008) Representation of geometric borders in the entorhinal cortex. Science 322(5909):1865–8.

Stewart S, Jeewajee A, Wills TJ, Burgess N, Lever C. (2014) Boundary coding in the rat subiculum. Philos Trans R Soc Lond B Biol Sci. 369(1635):20120514.

Taube JS, Muller RU, Ranck JB. Jr. (1990) Head-direction cells recorded from the postsubiculum in freely moving rats. I. Description and quantitative analysis. J Neurosci. 10(2):420–35.

Whitlock, J. R., Pfuhl, G., Dagslott, N., Moser, M.-B., & Moser, E. I. (2012) Functional split between parietal and entorhinal cortices in the rat. Neuron, 73(4), 789–802.

Wilson MA, McNaughton BL. (1993) Dynamics of the hippocampal ensemble code for space. Science 261(5124):1055–8.

Witter MP.(2006) Connections of the subiculum of the rat: topography in relation to columnar and laminar organization. Behav. Brain Res. 174:251–64.

Wood, E. R., Dudchenko, P. A., Robitsek, R. J., & Eichenbaum, H. (2000). Hippocampal neurons encode information about different types of memory episodes occurring in the same location. Neuron, 27(3), 623–33.

